# Characterization of an NADPH-dependent 17α-hydroxysteroid dehydrogenase encoded by the *desF* gene from the gut bacterium *Clostridium scindens* VPI 12708

**DOI:** 10.64898/2025.12.17.694922

**Authors:** Taojun Wang, Briawna Binion, João M.P. Alves, Jason M. Ridlon

## Abstract

Epitestosterone (epiT) is the isomer of the androgen testosterone. Historically, the role of epiT has remained unclear. Recently, it has been reported that epiT promotes AR-dependent prostate cancer cell proliferation. The gut bacterium *Clostridium scindens* VPI 12708 converts androstenedione (AD) to epiT. The bacterial enzymatic pathways involved in epiT formation have been reported, where the *desF* gene that encodes 17α-hydroxysteroid dehydrogenase converts AD to epiT using NADPH as a cofactor. In this study, we quantitatively characterized DesF kinetic parameters and substrate specificity. The results revealed that the optimal pH for the reductive reaction is 7.0, and for the oxidative reaction it is 7.5 and 8.0. The kinetic analysis showed that for the reductive reaction, the *K*_*M*_ was 8.67 ± 2.04 µM and the *V*_*max*_ was 1.95 ± 0.11 µM min^-1^; for the oxidative direction, the *K*_*M*_ was 27.17 ± 3.56 µM and the *V*_*max*_ was 2.18 ± 0.08 µM min^-1^. Moreover, the substrate specificity analysis revealed that 11-keto-AD is the most favourable substrate for DesF, and the 17-keto group of 11-keto-AD can be converted to the 17α-hydroxy group. These results are a significant advance in understanding epiT formation by the gut microbiome.

## 1. Introduction

Previous studies in the 1950s by Wade and Nabarro [1, 2] established the metabolism of cortisone by the gut microbiota in humans. Rectal infusion of cortisol resulted in elevated urinary excretion of 17-keto-steroids which was ablated by neomycin treatment [2]. Decades later Cerone-McLernon et al. [3] determined the major metabolites, including 21-deoxycortisol, tetrahydrocortisol, as well as the side-chain cleavage products of cortisol including 17-keto derivatives and 11β,17β-dihydroandrosterone. The first gut bacterial isolate with steroid-17,20-desmolase and steroid 20α-HSDH activity was reported in 1984 [4]. Originally named *Clostridium* sp. strain 19 [5], it was later classified as *Clostridium “scindens”* ATCC 35704^T^, becoming the type strain.

In 2013, we reported the identification of the *desABCD* gene cluster in *C. scindens* ATCC 35704 involved in both steroid-17,20-desmolase (*desAB*), NADH-dependent 20α-hydroxysteroid dehydrogenase (*desC*), and a predicted cortisol transporter (*desD*) [6]. Identification of *desAB* in *C. scindens* ATCC 35704 led to the identification of a gene cluster in *Butyricicoccus desmolans* and *Clostridium cadaveris*, which included the *desE* gene that encodes NADH-dependent 20β-HSDH [7]. We went on to solve the 2.0Å crystal structure of DesE from *Bifidobacterium adolescentis* [8]. Most recently, we showed that the combined culturing of *Clostridium scindens* ATCC 35704 (*des*AB) and *C. scindens* VPI 12708 (17α-HSDH activity) was capable of converting 11-deoxycortisol to androstenedione (AD) and epitestosterone (epiT) [9]. De Prada and colleagues (1994) previously demonstrated that 17α-HSDH activity in *C. scindens* VPI 12708 was inducible by the addition of either AD or cholic acid to the culture medium during the initiation of growth [10]. By comparing transcriptomes between 11β-hydroxy-androstenedione (11OHAD) induced vs. uninduced cells of *C. scindens* VPI 12708, we identified a single gene significantly upregulated that was annotated in the short chain dehydrogenase/reductase superfamily [9], of which many HSDHs are found [11]. We confirmed bidirectional NADP(H) conversion between AD and epiT by the recombinant 17α-HSDH, and proposed the name *desF* for this gene [9].

Here, we characterize recombinant DesF to determine oligomeric form, pH optima, kinetic constants, steroid substrate specificity and the phylogeny of DesF with other functionally confirmed HSDH sequences. This study is a significant advance in understanding the enzymatic basis by which the gut microbiome contributes to the formation of the potentially novel androgens, such as epiT.

## 2. Materials and Methods

### 2.1 Bacteria and chemicals

*Clostridium scindens* VPI 12708 was derived from in-house 30% glycerol stock cultures and cultivated in anaerobic Trypticase Soy Broth (TSB) at 37 °C. *Escherichia coli* DH5α competent cells (Thermo Scientific, Carlsbad, CA, USA) and *E. coli* BL-21 CodonPlus (DE3) RIPL (Agilent, Cedar Creek, Texas, USA) were purchased and incubated with Lysogeny broth (LB).

Steroids (commercial sources) used in this study included: 4-androstene-3,17-dione (Androstenedione, Steraloids, Newport, RI, USA); 4-androsten-17α-ol-3-one (Epitestosterone, Steraloids); 5α-androstan-3, 17-dione (5α-AD, Steraloids); 4-androsten-3, 11, 17-trione (11KAD, Steraloids); 4-androsten-11β-ol-3, 17-dione (11OHAD, Steraloids); 5α-androstan-17α-ol-3-one (5α-epitestosterone, Steraloids); 1,4-androstadiene-3,11,17-trione (AT, Sigma); All of the steroids were dissolved in DMSO for the usage.

### 2.2 Bacterial media preparation

The TSB (BBL) was prepared as instructed with the addition of 5 g yeast extract, 1 g L-cysteine, 1 mg resazurin and 40 mL salt solution (1 L; 0.25 g CaCl_2_•2H_2_O, 0.5 g MgSO_4_•7H_2_O, 1 g K_2_HPO_4_, 1 g KH_2_PO_4_, 10 g NaHCO_3_, 2 g NaCl). LB was purchased and prepared as instructed. LB Agar plates were prepared by adding 1.5% (w/w) agar to the LB broth.

### 2.3 Scanning electron microscopy (SEM)

*Clostridium scindens* VPI 12708 was incubated in TSB. Cultures (500 µL) in log phase were mixed with Karnovsky’s fixative (500 µL, 2% glutaraldehyde and 2.5% paraformaldehyde), vortexed, and then centrifuged at 4,000 x *g* for 5 min at 4°C to fix the bacterial isolates. Supernatant was discarded, and the pellet was washed 3 times and then sent to Materials Research Laboratory Central Research Facilities (University of Illinois at Urbana-Champaign, Urbana, Illinois, United States) for imaging with the Hitachi S-4800 high-resolution scanning electron microscope.

### 2.4 DesF heterologous expression and purification

DesF heterologous expression and purification were as described by Wang et al [9]. Briefly, the target inserts were amplified and inserted into the pET-51b(+) plasmid (Novagen, San Diego, CA, USA). Subsequently, the recombinant plasmid was selected by transforming it into *E. coli* DH5α competent cells. The target plasmid was transformed into *E. coli* BL-21 CodonPlus (DE3) RIPL. The *E. coli* BL-21 CodonPlus (DE3) RIPL was incubated overnight with 0.1 mM IPTG added to induce protein production. The microbial pellets were collected and lysed with 750 μL of lysozyme (1 mg/mL) and 10 μL of benzonase nuclease (Sigma). Afterwards, microbial pellets were physically lysed by passing through a French pressure twice. The recombinant proteins in the soluble fraction were purified using Strep-Tactin resins (IBA lifesciences) according to the manufacturer’s instructions. The purified proteins were assessed by sodium dodecyl sulfate-polyacrylamide gel electrophoresis (SDS-PAGE). Protein concentrations were measured by a Nanodrop 2000c spectrophotometer based on their extinction coefficients and molecular weights.

### 2.5 Proteomics

Samples of affinity purified DesF were digested with Trypsin (Thermo) using a CEM Discover Microwave Digestor (Matthews, NC) at 55°C for 30 min. The digested peptides were extracted, lyophilized, and cleaned up using stage-tips (Rappsilber, Mann et al. 2007). LC/MS was performed using a Thermo Fusion Orbitrap mass spectrometer in conjunction with a Thermo RSLC 3000 nano-UPLC. The column used was a Thermo PepMap C-18 (0.75 mm x 25 cm) operating at 300 nL/min and 40°C. The gradient was from 1% to 35% acetonitrile + 0.1% formic acid over a 45 min time interval. The mass spectrometer was operating in positive mode using data dependent acquisition method and collision induced dissociation for fragmentation at 35% energy. The raw data were analyzed using Mascot (Matrix Science, London, UK), and searches were made using a database consisting of the DesF sequences. FDR (False Data Discovery Rate) using a decoy reversed sequence database was at 1%.

### 2.6 Enzyme assays

Purified recombinant DesF activities were determined by mixing 10 nM enzyme, 50 μM substrate, and 200 μM cofactor (NADPH/NADP^+^) in phosphate-buffered saline. Samples were collected for metabolite extraction. Extracted samples were sent to the Mass Spectrometry Lab (University of Illinois at Urbana-Champaign, Urbana, Illinois, United States) for metabolite analysis using LC-MS.

### 2.7 Steroid extraction

The samples and ethyl acetate in a 1:2 ratio were thoroughly mixed by vortexing for 1 min. Next, the ethyl acetate layer was carefully collected and transferred to new tubes. The extraction process was repeated, and the collected top layers were evaporated with nitrogen gas and dissolved in 200 µL of methanol for LC-MS analysis.

### 2.8 Liquid chromatography-mass spectrometry (LC-MS)

Samples were sent to the Mass Spectrometry Lab (University of Illinois at Urbana-Champaign, Urbana, Illinois, USA) for metabolite analysis using liquid chromatography-mass spectrometry (LC-MS). LC-MS for all samples was done as described by Wang et al [9]. Briefly, a Waters Aquity UPLC coupled with a Waters Synapt G2-Si ESI MS (Waters Corp., Milford, MA, USA) was used for the sample analysis with a Waters Acquity UPLC BEH C18 column (1.7 μm particle size, 2.1 mm × 50 mm). The column temperature was maintained at 40°C with an injection volume of 0.5 µL. For gradient elution, 2 mobile phases were used: mobile phase A contained 95% water, 5% acetonitrile, and 0.1% formic acid; mobile phase B contained 95% acetonitrile, 5% water, and 0.1% formic acid. The flow rate was 0.5 mL/min. The LC eluents were introduced into the mass spectrometer equipped with electrospray ionization (ESI) with a positive ion mode for steroid analysis. Mass Lynx v4.1 (Waters) was used for generating chromatographs and mass spectrometry data analysis.

### 2.9 pH optimization

To determine the optimal pH of DesF in both oxidative and reductive directions, buffers with different pH values were applied. The buffers were as follows: 50 mM sodium citrate, 150 mM NaCl (pH 5.0 to 5.5); 50 mM sodium phosphate, 150 mM NaCl (pH 6.0 to 7.5); 50 mM Tris base, 150 mM NaCl (pH 8.0 to 9.0); 50 mM glycine, 150 mM NaCl (pH 9.5 to 10). A total of 200 µL buffer with 50 µM AD, 150 µM NADPH and 0.01 µM enzyme was used for oxidative reactions. A total of 200 µL buffer with 50 µM epiT, 150 µM NADP^+^ and 0.01 µM enzyme was used for the reductive reactions. Absorbance was measured over time at 340 nm using the BioTek Epoch 2 Microplate Spectrophotometer (Agilent, Santa Clara, CA, USA).

### 2.10 Enzyme Kinetics

Kinetic was determined at the optimal pH values. A total of 200 µL buffer (pH = 7.0, 50 mM sodium phosphate, 150 mM NaCl) with 150 µM NADPH, 0.01 µM enzyme and 0, 1, 2, 5, 10, 20, 50, 100, 200, 300 µM AD, respectively, was used for oxidative reactions. A total of 200 µL buffer (pH = 7.5, 50 mM sodium phosphate, 150 mM NaCl) with 150 µM NADP^+^, 0.01 µM enzyme and 0, 1, 2, 5, 10, 20, 50, 100, 200, 300 µM epiT, respectively, was used for the reductive reactions. Absorbance was measured over time at 340 nm using the BioTek Epoch 2 Microplate Spectrophotometer (Agilent). The kinetic analysis was done by fitting the data to the Michaelis-Menten equation using the *drc* v3.0-1 package [12] with R version 4.3.0 [13].

### 2.11 Phylogenetics

Maximum-likelihood phylogenetic analysis was performed on selected HSDH protein sequences from a variety of taxa of interest using RaxML v. 8.2.12 [14]. Sequences were based on selections used in prior analysis [15, 16]. The best-fitting substitution model (LG with empirical residue frequencies) was chosen automatically by the program. Site substitution rate heterogeneity was modeled using gamma distribution-approximated categories. Statistical robustness of the inferred nodes was evaluated by boostrap, with 700 pseudoreplicates being performed (the amount was automatically determined by the software). The tree display and annotation were done by an online tool iTOL [17].

## 3. Results

### 3.1 Expression and purification of recombinant DesF

*Clostridium scindens* VPI 12708 (**Figure 1A**) converts AD to epiT by 17α-hydroxysteroid dehydrogenase (HSDH). We previously cloned the *desF* gene encoding 17α-HSDH into pET-51b(+) and verified the integrity of the DNA sequence (Wang, Ahmad et al. 2025). The recombinant DesF (rDesF) was expressed in *E. coli* BL-21 CodonPlus (DE3) RIPL and purified to electrophoretic homogeneity (**Figure 1B**), and the amino acid sequence of the purified recombinant DesF (rDesF) was verified by proteomic analysis (**Supplementary Table 1**). The molecular mass was determined to be 30.11 KDa by mass spectrometry. We have confirmed that DesF can convert AD to epiT (reductive reaction) using NADPH as cofactor and DesF can convert epiT to AD (oxidative reaction) using NADP^+^ as cofactor (**Figure 1C**).

**Table 1.** Michaelis-Menten enzyme kinetic parameters.

| **Michaelis-Menten** |  |  |  |  |
| --- | --- | --- | --- | --- |
| Substrates | V<sub>max</sub> (μM/min) | K<sub>m</sub> (μM) | K<sub>cat</sub> (min<sup>-1</sup>) | K<sub>cat</sub>/K<sub>m</sub> (μM<sup>-1</sup>·min<sup>-1</sup>) |
| Androstenedione | 1.95 ± 0.11 | 8.67 ± 2.04 | 194.54 ± 10.54 | 2.23 ± 0.54 |
| Epitestosterone | 2.18 ± 0.08 | 27.17 ± 3.56 | 217.71 ± 7.97 | 2.23 ± 0.30 |

**Figure 1.**
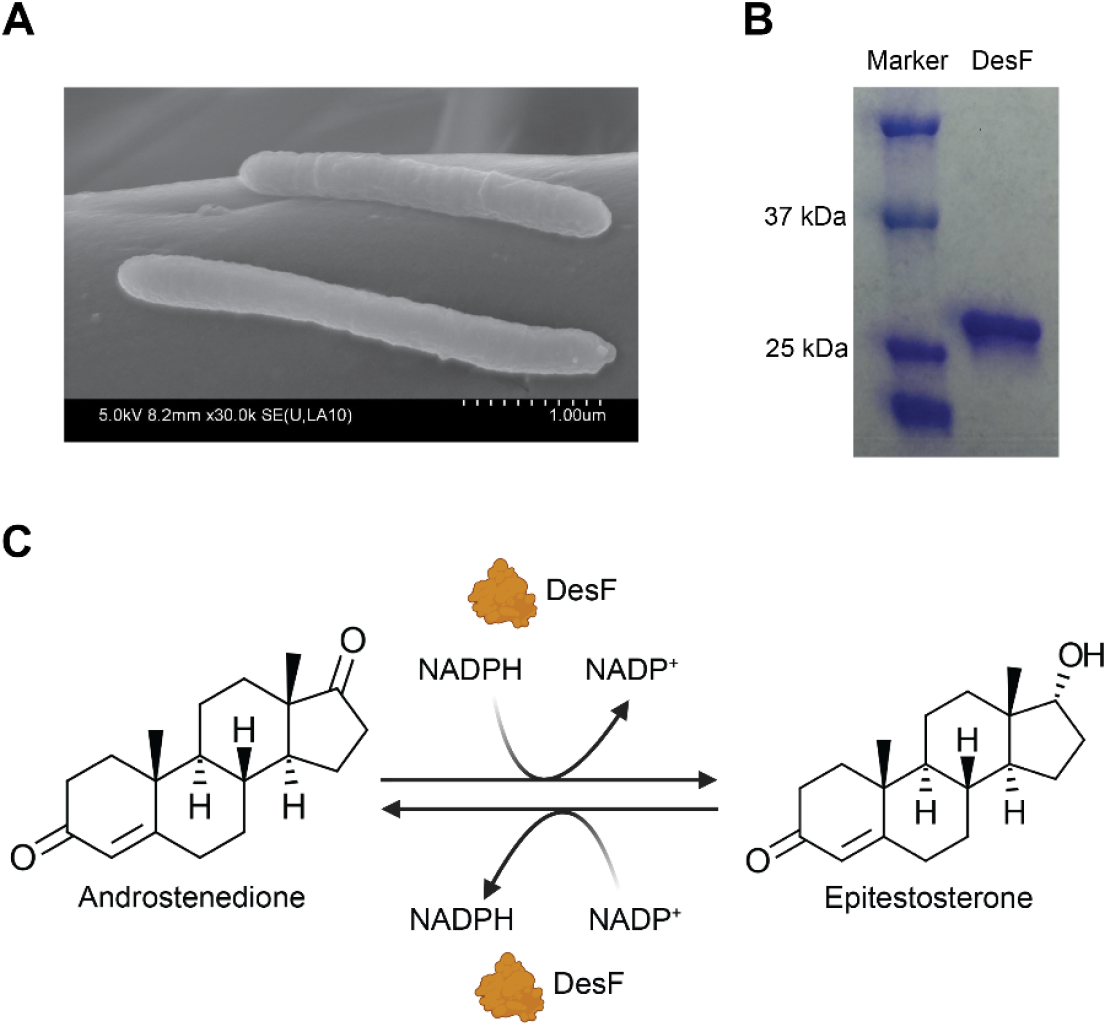
DesF (17α-hydroxysteroid dehydrogenase) from *Clostridium scindens* VPI 12708 converts androstenedione to epitestosterone, vice versa. (**A)** SEM image of *C. scindens* VPI 12708. (**B**) SDS-PAGE of recombinant DesF. (**C**) Reactions of DesF using androstenedione or epitestosterone as substrates, respectively.

### 3.2 DesF optimal pH and kinetics

We first optimized the pH for the reductive and oxidative reactions by combining 50 µM steroid substrate, 150 µM NADPH and 0.01 µM enzyme in a series of buffers from pH 5.5 to 8.5 in the reductive direction and pH 6.0 to 9.0 in the oxidative direction (**Figure 2A**). In the reductive direction, relative activity climbs from 71.06 ± 1.72% relative activity at pH 5.5 to an optimum pH at 7.0 before steadily declining to 17.45 ± 1.88% relative activity at pH 8.5. In the oxidative direction, relative activity increased from 26.64 ± 2.09% at pH 6.0 to an optimum between pH 7.5 and 8.0 before descending to 43.05 ± 2.03% relative activity at pH 9.0 (**Figure 2B**).

**Figure 2.**
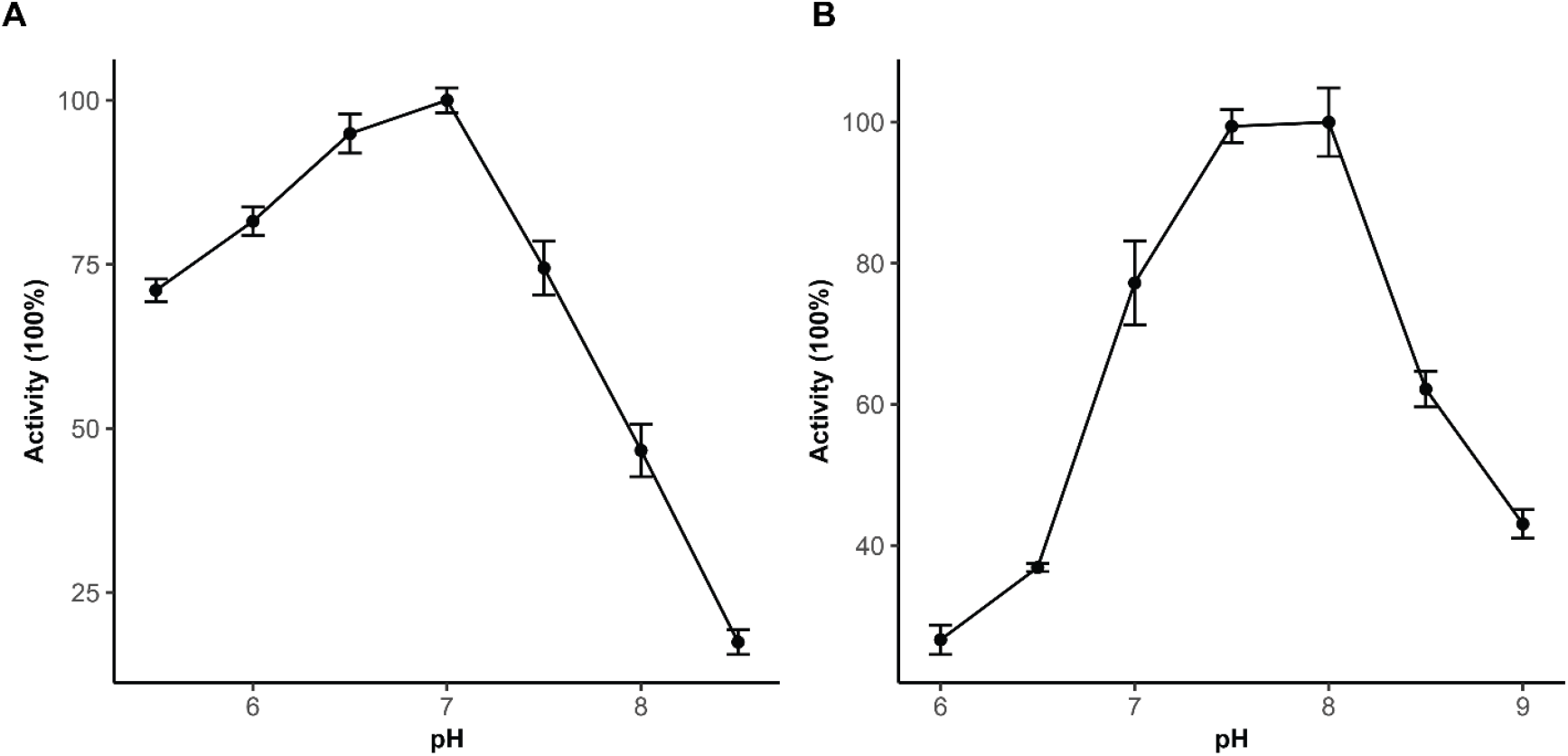
Determination of optimal pH of DesF using androstenedione (**A**) or epitestosterone (**B**) as substrate.

For kinetic analysis, we chose pH 7.0 for the reductive direction using AD as substrate (**Figure 3A**) and pH 7.5 for the oxidative direction using epiT as substrate (**Figure 3B**). In the reductive direction, we varied AD from 0 to 300 µM while holding NADPH constant at 150 µM. The *K*_*M*_ was 8.67 ± 2.04 µM and the *V*_*max*_ was 1.95 ± 0.11 µM min^-1^. The turnover number (*k*_*cat*_) was determined to be 194.54 ± 10.54 min^-1^ and the catalytic efficiency (*k*_*cat*_/*K*_*M*_) was determined to be 2.23 ± 0.54 µM^-1^ min^-1^. For the oxidative direction, the *K*_*M*_ was 27.17 ± 3.56 µM and the *V*_*max*_ was 2.18 ± 0.08 µM min^-1^. The turnover number (*k*_*cat*_) was determined to be 217.71 ± 7.67 min^-1^ and a catalytic efficiency (*k*_*cat*_/*K*_*M*_) of 2.23 ± 0.03 µM^-1^ min^-1^ (**Table 1**).

**Figure 3.**
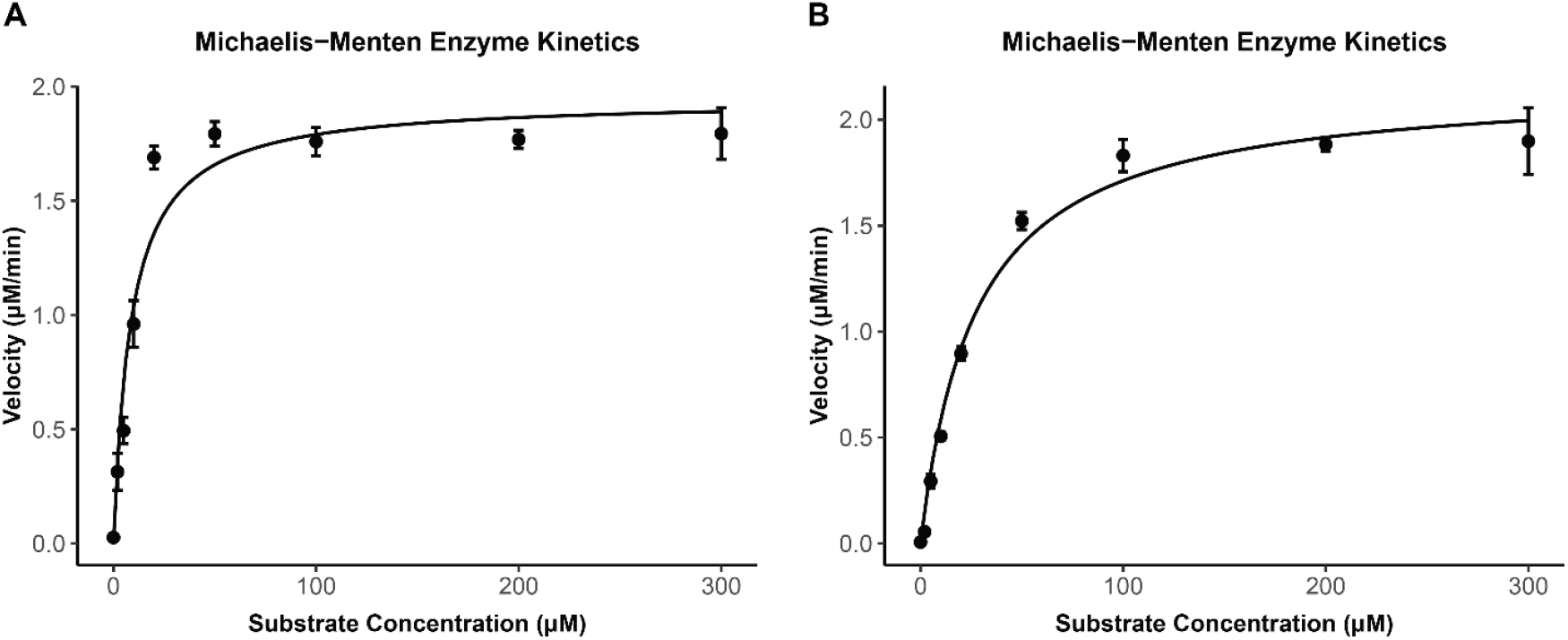
Michaelis-Menten enzyme kinetics of DesF using androstenedione (**A**) or epitestosterone (**B**) as substrate.

### 3.3 DesF substrate specificity

We then performed substrate specificity studies with rDesF. Previously, we confirmed stereospecificity for substrates with a 17α-hydroxy group in the oxidative direction [9]. We therefore excluded 17β-hydroxy steroids such as testosterone. In the reductive direction, we tested AD (androst-4-ene-3,17-dione; AD), androstanedione (5α-androstane-3,17-dione; 5α-AD), 11β-hydroxyandrostenedione (11β-OHAD), 11keto-androstenedione (11K-AD) and the prednisone side-chain cleavage product, 1,4-androstadien-3,11,17-trione (AT). We set AD at 100% relative activity, but we found that 11K-AD was the best substrate at 154.79 ± 9.59 % relative activity (**Table 2**). Interestingly, reduction of the 11keto-group in the form of 11β-OHAD reduced activity to 39.36 ± 1.60 %. Modifications to the A-ring led to a reduction to 46.45 ± 2.51 % relative activity in the case of 5α-AD and 51.95 ± 1.63 % relative activity in the case of AT (**Table 2**). In the oxidative direction, 5α-epitestoterone showed 72.63 ± 5.58 % activity relative to epiT. We then confirmed product formation by LC-MS (**Figure 4**).

**Table 2.**
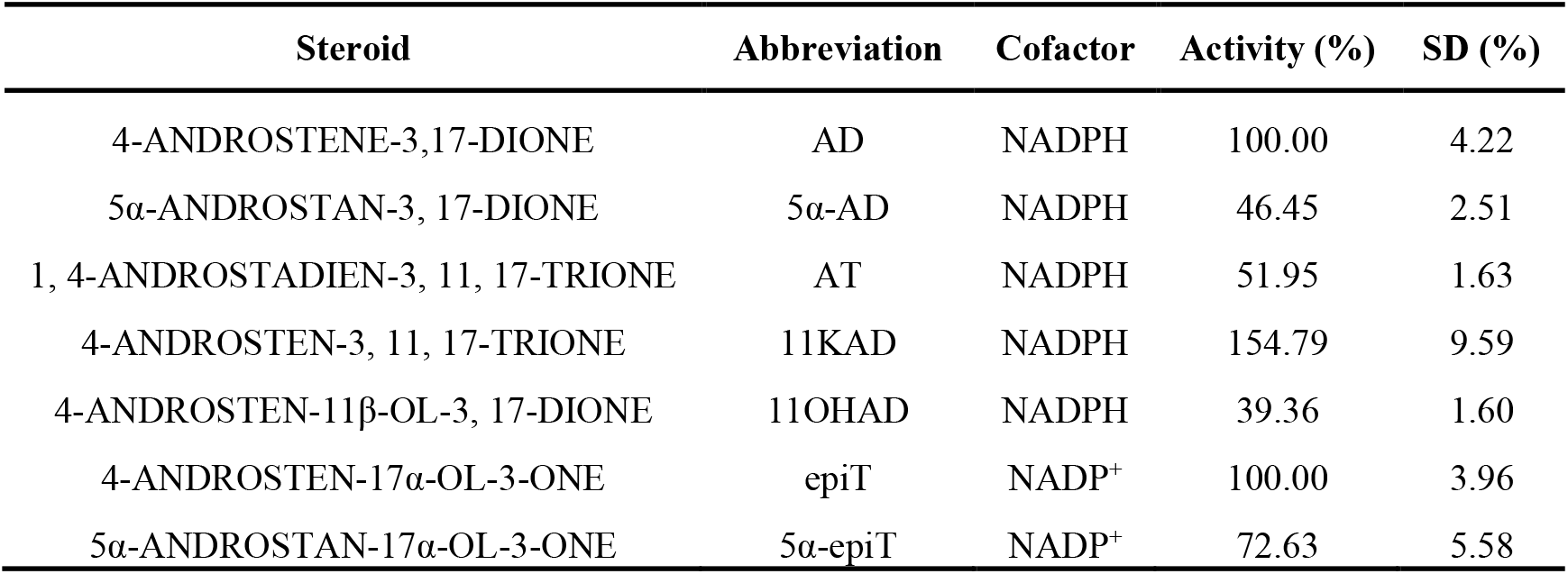
Substrate specificity of DesF.

**Figure 4.**
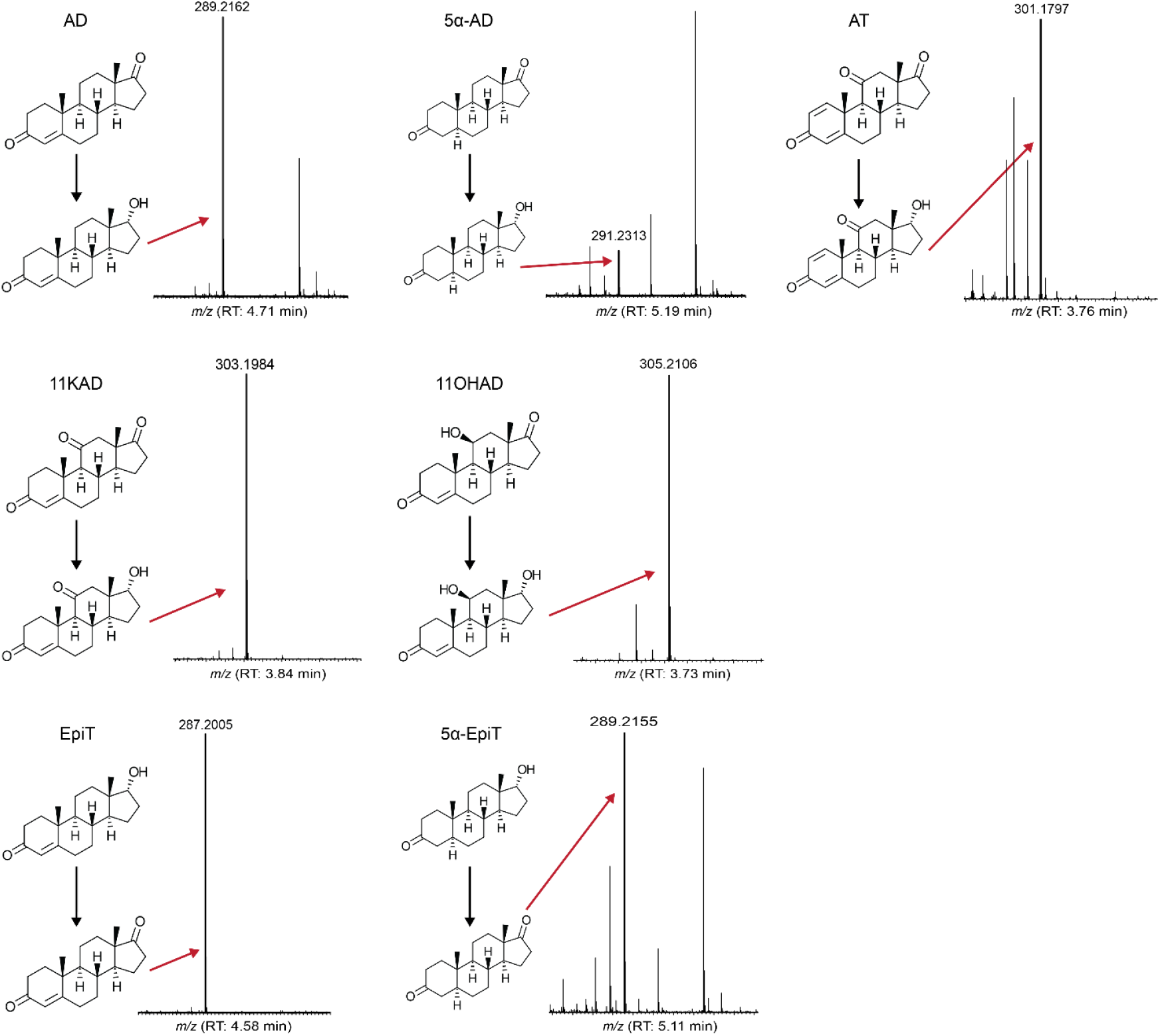
Confirmation of substrate specificity of DesF using LC-MS.

### 3.4 Phylogenetics of DesF

The phylogenetic relationships between DesF and stereospecific HSDHs from bacteria and vertebrate were investigated (**Figure 5**). In general, most of the HSDHs are clustered together according to their known stereospecificity, and the bacterial HSDHs are largely separated from their vertebrate counterparts. This is true of bacterial 7α-HSDHs from gut Bacillota and Bacteroidota, gut and urinary tract 20β-HSDHs (DesE) [7], bacterial 12β-HSDHs [16], human 3β-HSD/Δ^4,5^-isomerases. DesC (WP_004606450.1) from *Clostridium scindens* ATCC 35704 [6], encoding 20α-HSDH, can convert cortisol to 20α-dihydrocortisol, was separated from 20β-HSDH. To our knowledge, DesF is the only reported host-associated bacterial 17α-HSDH. DesF is clustered with 17β-HSDH type 10 (AAC15902.1) [18] and 3β-HSDH (WP_015759877.1) from *E. lenta*. Of interest is that AAC15902.1 is a protein from humans, indicating that these SDR family enzymes might share an early origin and have a high degree of conservation in evolution. DesG is a 17β-HSDH that is part of a branch that shares common ancestry with DesF. DesG clustered with WP_076388097.1 (17β-HSDH) from *Vaginimicrobium propionicum* and WP_015760525.1 (3β-HSDH) from *E. lenta*.

**Figure 5.**
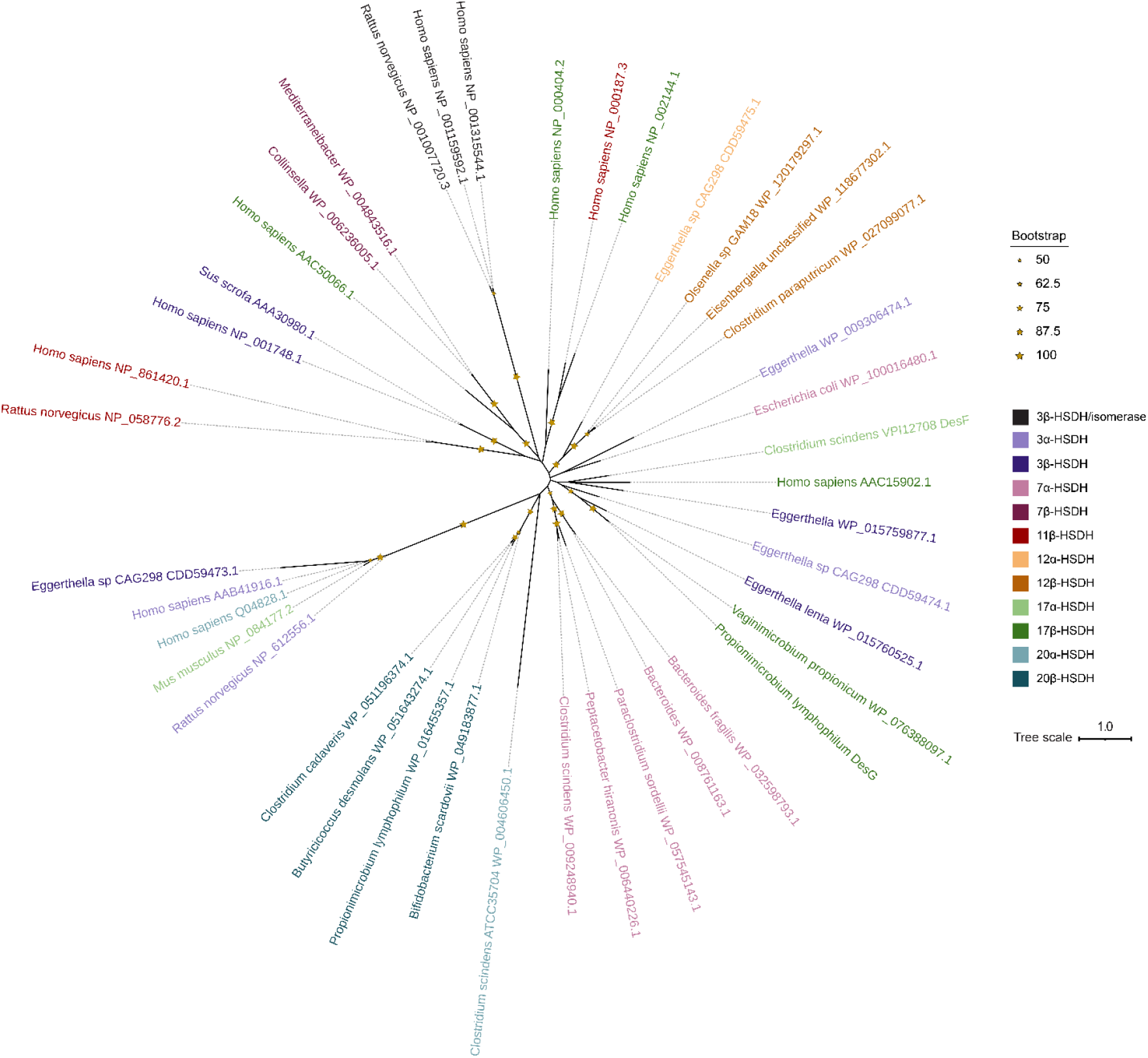
Maximum-likelihood phylogenetic analysis of HSDHs. Clusters are colored by function as displayed in the legend. Bootstrap values are displayed by the size of stars.

## 4. Discussion

Over thirty years ago, *Eubacterium* sp. strain VPI 12708 (now *C. scindens* VPI 12708) was reported to convert AD to epiT [10]. N-terminal sequencing of the native semi-purified fraction containing 17α-HSDH activity in that study suggested a novel 42 kDa disulfide reductase catalyzes the reaction [10]. Recently, we applied RNA-Seq to discover a steroid-inducible gene (*desF*) encoding a 30.11 kDa short chain dehydrogenase/reductase family protein with 17α-HSDH activity [9]. The current study expands upon our understanding of microbial steroid metabolism by DesF through demonstrating that steroids of adrenal origin (e.g. 11K-AD, 11β-OHAD) are substrates. To our knowledge, the resultant metabolites, 11K-epiT and 11β-OH-epiT have not been reported and may represent unique microbial metabolites whose physiological effects are unknown. Indeed, recent work has challenged the widely held notion that epiT acts as an “anti-androgen” and suggests instead that derivatives of epiT act as ligands for the androgen receptor [9, 19].

Interestingly, recent pangenome analysis of *C. scindens* resulted in strains grouping into two distinct phylogenetic clusters with differences in average nucleotide identity sufficient to categorize these clusters as distinct species [20]. Indeed, the strains that have been archetypes in the study of bile acid and steroid metabolism, *C. scindens* VPI 12708 and *C. scindens* ATCC 35704, separated into the two distinct clusters. The “VPI 12708” group, with only a few exceptions, encoded DesF which was not found in strains in the “ATCC 35704” cluster. Conversely, the “ATCC 35704” cluster was characterized by encoding *desABCD* gene clusters. *C. scindens* S077 (VPI 12708 cluster) was the only exception to this trend in harboring *desABCD* and *desF* genes and we demonstrated the ability of this strain to convert 11-deoxy-cortisol to epiT [9]. This underscores the importance of recognizing and taking into account strain variation in steroid metabolic capacity in the microbiome.

The role of circulating epiT, as well as its route of biosynthesis by host enzymes, remain unclear in humans. With few exceptions [19, 21], epiT derivatives have traditionally been thought to be “antiandrogens” with relatively high circulating concentrations on par with testosterone in some men [22]. We recently determined that epiT causes AR-dependent proliferation in LNCaP prostate cancer cell line (expresses mutant AR) to a significantly greater extent than testosterone. Proliferation caused by epiT treatment was also observed in VCaP prostate cancer cells that express wild type AR [9]. Additionally, the side-chain cleavage product of prednisone, 1,4-androstenedione (AT), when converted to epiAT by *C. scindens* VPI 12708 caused significant growth of LNCaP cells in an AR-dependent manner [9]. In this study, we further showed that DesF converted AT to epiAT using NADPH as a cofactor. Importantly, the inhibitor of host steroid-17,20-desmolase (CYP17A1), abiraterone, did not inhibit bacterial steroid-17,20-desmolase in *C. scindens* ATCC 35704. Moreover, we observed that DesF convert 5α-reduced AD to 5α-reduced epiT, and the recent report indicated that 5α-reduced epiT (17α-hydroxy-5α-androstan-3-one; epiDHT) agonizes nuclear AR [19].

Taken in this context, the distinction between these clades of *C. scindens* is expected to be clinically important. During treatment for castration resistant prostate cancer, the drug abiraterone acetate (AA) blocks the formation of adrenal androgens and their precursors (e.g. DHEA, AD, 11OHAD) as well as the glucocorticoid cortisol (but not corticosterone) by inhibiting host steroid-17,20-desmolase CYP17A1. The inhibition of cortisol biosynthesis requires replacement of cortisol with prednisone (P) [23]. Those responding to AA/P treatment are defined as maintaining stable blood prostate specific antigen (PSA) levels corresponding to low levels of circulating androgens. Inevitably, men undergoing AA/P treatment will begin experiencing rising blood PSA, indicating the return of rising circulating androgens and disease progression. Determining the source of renewed androgen biosynthesis is critical to improving current androgen deprivation therapy. We developed quantitative PCR (qPCR) primers to target the *desF* gene in fecal samples from hormone sensitive prostate cancer (HSPC) versus those undergoing treatment for advanced prostate cancer with (AA/P) to determine whether fecal *desF* gene levels correlate to response to AA/P treatment [9]. Indeed, both absolute *desF* gene copy number and normalized *desF* gene abundance to total 16S rRNA were significantly different between HSPC subjects and those progressing on AA/P. The gut microbiome of some men may harbor *desF* only, in which case endogenous AD may be converted to epiT potentially resulting in some elevation of PSA due to androgen signaling. However, if the same individual harbored both *desAB* and *desF*, both prednisone and AD would be expected to contribute to a more substantial rise in blood PSA due to elevated circulating androgens. Taken together, strains of *C. scindens* may be a source of androgens (e.g. epiAT) from biotransformation of prednisone in the context of advanced prostate cancer, and the enzymology of the steroid-17,20-desmolase pathway, including *desF*, is of potential biomedical importance [24, 25].

